# Divide and conquer: a viral bifunctional RNA utilizes both RNA and protein to modulate pro-tumourigenic pathways

**DOI:** 10.64898/2026.10.26.762023

**Authors:** Maxwell P. Bui-Marinos, Jennifer A. Corcoran

**Affiliations:** Microbiology, Immunology and Infectious Diseases Department Arnie Charbonneau Cancer Institute, Robson DNA Science Centre, Snyder Institute for Chronic Diseases Cumming School of Medicine, University of Calgary, Calgary, Alberta, Canada

## Abstract

Kaposi’s sarcoma-associated herpesvirus (KSHV) is an oncogenic herpesvirus that causes the endothelial cell neoplasm Kaposi’s sarcoma (KS) and other B-cell malignancies. The viral transcript *kaposin* is largely composed of GC-rich direct repeats (DRs) and is expressed during all phases of KSHV replication. *Kaposin* is polycistronic, encoding for at least 3 kaposin (Kap) proteins; of these, KapB is the most abundant and well-studied. Recent work from our lab discovered that *kaposin* is found in nuclear speckles (NSs) and that DRs are required for NS retention. During latency, *kaposin* is predominantly in NSs and is not exported or translated into KapB. During viral reactivation, abundant *kaposin* correlates with *kaposin* escape from NSs and KapB protein production. Thus, although *kaposin* is a latent transcript, KapB protein production occurs only during lytic reactivation. The spatiotemporal regulation of *kaposin* translation by NS retention during KSHV latency likewise suggests that these two biomolecules may have distinct functions. Because historically *kaposin* was viewed as a protein-coding mRNA, overexpression studies attributed any function to KapB even though both *kaposin* and KapB were produced in these systems. Armed with the knowledge that *kaposin* is a bifunctional RNA which derives function from both the transcript and the protein(s) it encodes, it remains unclear whether prior phenotypes should be attributed to *kaposin* or KapB. In this study, we developed a reductionist overexpression system to uncouple the effects of these biomolecules. To mimic *kaposin* during viral latency, we used a construct capable of forming *kaposin* NSs that does not translate reported Kap proteins (ΔStarts). To mimic late lytic reactivation, we used a recoded construct, AltB, that does not form *kaposin* NSs, is readily exported from the nucleus, and translates abundant KapB. We found that KapB protein induced actin stress fibre formation, cell enlargement, increased total levels of Yes-associated protein (YAP) and increased YAP-target gene transcription including pro-growth and pro-inflammatory transcripts. Conversely, *kaposin* RNA induced cell autonomous p-body disassembly with an overall reduction in pro-growth and pro-inflammatory transcripts, suggesting *kaposin* might sustain cell survival while dampening immune signalling. Taken together, we demonstrate that *kaposin* RNA and KapB protein have distinct and sometimes opposing effects on cellular growth and inflammatory signalling pathways, highlighting the ability of a viral bifunctional RNA to use both its RNA and protein products to mediate viral tumourigenesis.

## INTRODUCTION

Kaposi’s sarcoma-associated herpesvirus (KSHV) is a large DNA oncovirus and known etiological agent of the endothelial cell neoplasm, Kaposi’s sarcoma (KS), two B-cell lymphoproliferative diseases, primary effusion lymphoma (PEL) and multicentric Castleman’s disease (MCD), and KSHV inflammatory cytokine syndrome (KICS) [1–4]. Like all herpesviruses, KSHV causes a lifelong infection in its human host, cycling between a persistent, latent infection and lytic replication. In latency, viral gene expression is restricted to only a subset of genes, the viral latency locus, allowing latently infected cells to maintain the viral genome while simultaneously evading the immune system [5, 6]. The endothelial cells (ECs) in KS lesions adopt a spindle-like morphology, and early studies using a riboprobe targeting the KSHV latency-associated T0.7 RNA transcript, now primarily known as *kaposin*, found that ECs are predominantly latently infected [7, 8]. Kaposin was identified as a transforming gene [9], is expressed throughout the viral life cycle [10–12], and *kaposin* RNA is consistently and abundantly detected in all stages of KS lesion development [7, 8, 13–15]. These collectively imply that the kaposin gene locus and its products are important contributors to viral tumorigenesis, yet the complex nature of kaposin leaves it understudied relative to other latency genes.

The *kaposin* RNA transcript contains two, GC-rich and highly repetitive regions termed direct repeat 1 (DR1) and direct repeat 2 (DR2), followed by the K12 open reading frame (ORF; *Fig. 1A*) [10]. DR1 and DR2 are each comprised of a unique set of 23 nucleotide repeats, reiterated a variable number of times depending on the viral isolate [16]. In this current study, the JSC-1 isolate derived from the KSHV-infected PEL cell line is used; it contains 27 copies of DR1 and 21 copies of DR2 [17, 18]. *Kaposin* RNA is polycistronic and predicted to encode for at least three proteins: Kaposin (Kap) A, KapB, and KapC (**Fig. 1A**) [10]. KapA translation is initiated from a canonical AUG start codon, capturing ORF K12, and is predicted to be a small transmembrane protein [10, 19]. KapC is translated in the same frame as KapA using one of two upstream, non-canonical CUG start codons, capturing kaposin amino acid direct repeats (Kap DRs) [10] (DR1 x 9; DR2 x 7). KapC also contains the K12 transmembrane domain at its C-terminal end. KapB translation is also initiated from one of two non-canonical start codons in a different frame than KapA and KapC. Due to differential stop codon usage prior to the K12 ORF, KapB is shorter and is comprised largely of DRs. Though *kaposin* is polycistronic, KapB is the dominantly expressed Kap protein [10] and has been the subject of several kaposin overexpression studies [20–29]; to date, KapB remains the most well characterized Kap protein.

**Figure 1.**
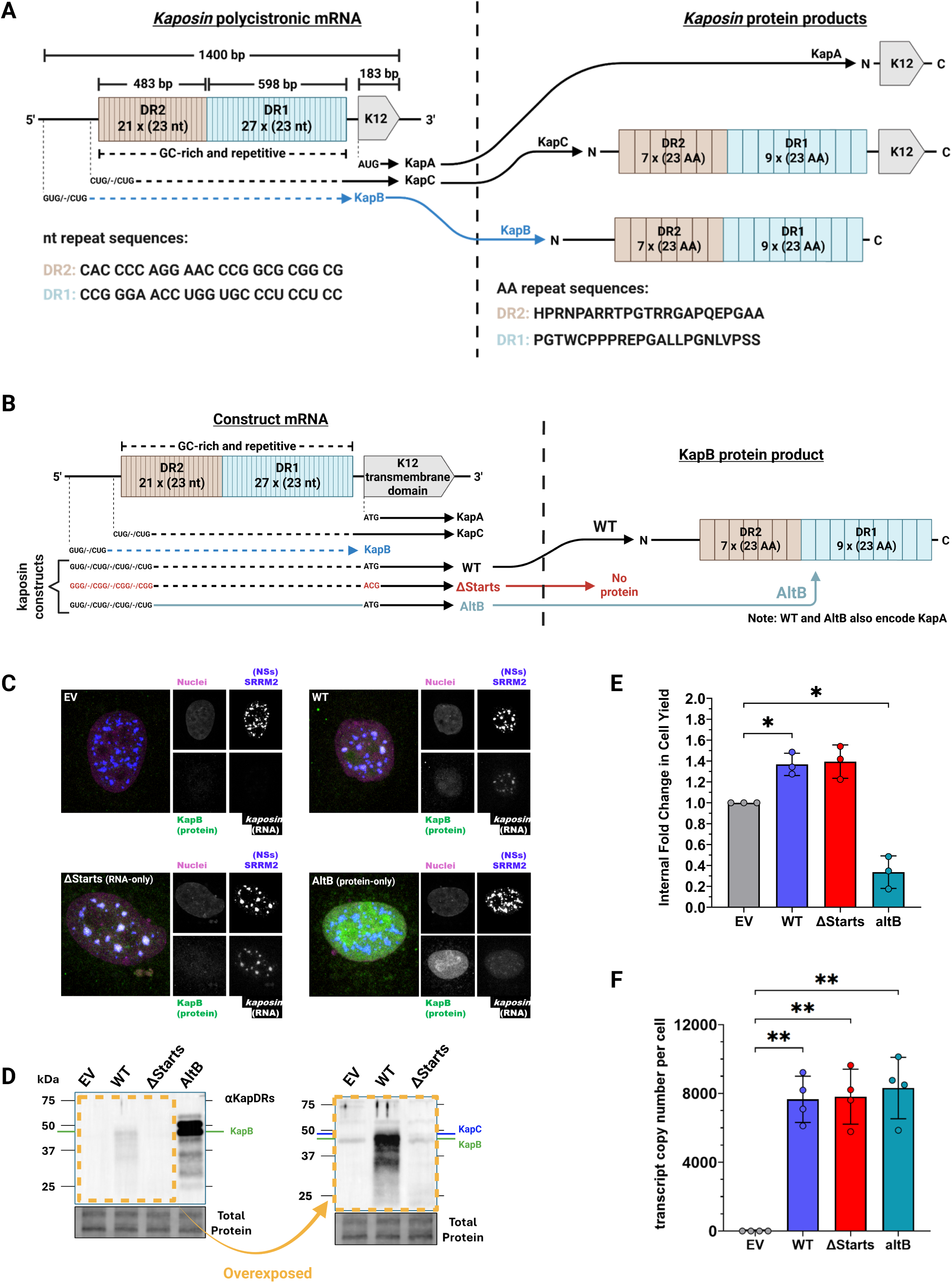
Design and characterization of kaposin constructs to uncouple kaposin RNA and KapB protein functions. **(A)** Schematic of the *kaposin* mRNA locus (left) and kaposin protein products (right). Under *kaposin* RNA, black and blue lines indicate native transcription of the *kaposin* locus into RNA, and blue reflects the fact that KapB is encoded in a different frame relative to KapA and KapC. Dashed lines indicate GC-rich and repetitive RNA regions; solid lines indicate non-GC-rich and non-repetitive RNA regions. “DR” = direct repeat, “nt” = nucleotide, and “AA” = amino acid. **(B)** Schematic of kaposin constructs, wild type (WT), ΔStarts, and AltB. Red colouring indicates nucleotide substitutions that eliminate start codons, and teal colouring indicates an alternatively coded RNA sequence. Dashed and solid lines as in **Fig 1A**. Only KapB translation is shown; KapA and KapC are translated in WT as in **Fig. 1A** and are not translated in ΔStarts while KapA is also translated in AltB. **(C)** A549 cells were transduced with lentiviruses encoding WT, ΔStarts, and AltB constructs alongside an empty vector (EV) control; post-selection, cells were fixed then immunostained using antibodies specific for NS-resident SRRM2 (false-coloured blue; AlexaFluor-488) and Kap DRs (false-coloured green; AlexaFluor-555) followed by RNA-FISH for *kaposin* RNA (false-coloured white; Quasar-670) and Hoechst stain for nuclei (false-coloured magenta). Photomicrographs were taken using an LSM 880 Airyscan confocal microscope and presented as maximum-intensity projection. One representative image set of three independent replicates (*n* = 3) is shown, and independent channels are shown in grayscale. Scale bar = 10 μm. **(D)** Protein lysates were harvested from A549 cells transduced with kaposin constructs and EV, post-selection, then used for immunoblotting using an antibody specific for Kap DRs. Total protein was used as a loading control. The green line indicates the position of KapB protein while the blue line indicates KapC. Since KapA lacks DRs, our antibody cannot detect the KapA protein. One representative blot of three independent replicates (*n* = 3) is shown. **(E)** A549 cells transduced to overexpress kaposin constructs and EV, post-selection, were counted using a hemocytometer. Kaposin construct cell counts were internally normalized to EV counts to calculate internal fold change in cell yield. Statistical analysis was conducted using a repeated-measures one-way ANOVA with a Tukey’s post-hoc (*n* = 3; mean ± SEM; * *p* < 0.05). **(F)** RNA was harvested from A549 cells transduced with kaposin constructs and EV, post-selection, and converted to cDNA which was then used as a template for RT-qPCR transcript copy analysis using a TaqMan assay specific for ORF K12 (KapA). Total *kaposin* transcripts across constructs were normalized by respective cell counts to calculate *kaposin* transcript copy number per cell. Statistical analysis was conducted as in **Fig. 1E** (*n* = 4; mean ± SEM; * *p* < 0.05, ** *p* < 0.01).

KapB is implicated in several processes relevant to KSHV pathogenesis. KapB localizes in the nucleus where it binds to and activates mitogen-activated protein kinase-activated protein kinase 2 (MAPKAPK2 or MK2) [20], a downstream effector of p38-MAPK signalling associated with inflammation and tumour progression (reviewed in [30–32]). This inhibited the degradation, or enhanced the stability, of mRNAs with AU-rich elements (AREs) in their 3’UTRs that encode for pro-inflammatory and pro-growth molecules like inteleukin-6 (IL-6) and granulocyte-macrophage colony-stimulating factor (GM-CSF) [20, 21]. KapB-mediated activation of MK2 caused Rho GTPase-dependent cytoskeletal rearrangements (reviewed in [33]) to induce cell spindling and actin stress fiber (SF) formation in ECs. These phenotypes were required for the disassembly of cytoplasmic RNA granules that harbor ARE-mRNAs called processing (p-)bodies (reviewed in [34]) [25, 26]. More recently, KapB-mediated actin remodeling was shown to activate the pro-growth, mechanosensitive transcriptional co-activator *Yes*-associated protein (YAP; reviewed in [35, 36]), YAP is also required for KapB-mediated p-body disassembly [28]. While this list of KapB-mediated phenotypes is non-exhaustive, all phenotypes attributed to Kap proteins have been in the context of ectopic overexpression. To understand the contribution of Kap proteins in infection, our lab generated a panel of recombinant KSHV mutants designed to lack Kap proteins individually or in combination [37]. However, all mutant viruses were impaired and maintained low viral episome copy number compared to wild type which was not restored when Kap proteins were supplied in *trans* [37]. This hinted towards *kaposin* RNA having a functional role independent of the Kap proteins, especially since all mutant viruses have a vastly altered *kaposin* RNA sequence.

Indeed, our lab recently discovered that *kaposin* RNA uses its GC-rich repeats to seed nuclear foci containing nuclear speckle (NS) machinery [38], such as serine/arginine repetitive matrix 2 (SRRM2) [39, 40]. NSs promote the RNA processing and expression of genes they are spatially proximal to [41, 42]; in KSHV infection, the formation of NSs seeded by *kaposin* (herein *kaposin* NSs) adjacent to the viral genome enhanced viral replication and gene expression [38]. Only viruses and overexpression constructs where *kaposin* contains intact DR1 and DR2 domains resulted in the formation of *kaposin* NSs. The nuclear retention of *kaposin* RNA makes it unavailable for translation during latency, and only later in the lytic cycle is *kaposin* exported and Kap proteins are detected [38], suggesting that NS retention provides spatiotemporal regulation of *kaposin* translation into Kap proteins. This discovery defines *kaposin* as a bifunctional RNA that utilizing both the protein it encodes (KapB) and the RNA (kaposin) for distinct purposes. Because all prior work used overexpression constructs that express both KapB and *kaposin*, it is now unclear if prior phenotypes attributed to KapB are caused by the RNA or the protein.

In this study, we sought to disentangle if established functions associated with KapB are elicited by *kaposin* RNA or KapB protein. To achieve this, we created a panel of “kaposin constructs” (**Fig. 1B**). Wild type (WT) encodes the full-length kaposin gene, producing GC-rich, repetitive *kaposin* RNA that encodes for all Kap proteins. In ΔStarts, all reported canonical and non-canonical start codons were eliminated to create an otherwise authentic *kaposin* RNA with GC-rich DR1 and DR2 regions intact that fails to translate KapA, B and C proteins. In AltB, we took advantage of codon degeneracy to produce a non-repetitive *kaposin* RNA that is less GC-rich (62% down from 74% GC-content), lacks DRs, and encodes the authentic KapB amino acid sequence and KapA, but fails to translate KapC. Using these kaposin constructs in lentivirus overexpression, we found that KapB induced actin stress fibre formation, increased cell size, and increased total YAP levels, while *kaposin* increased nuclear YAP. *Kaposin* expression disassembled p-bodies in a cell autonomous manner while KapB expression induced p-body disassembly using a paracrine mechanism. Consistent with this, KapB increased steady-state levels of growth and inflammatory transcripts whereas *kaposin* decreased these transcripts. Taken together, these findings demonstrate that *kaposin* RNA and KapB protein have distinct, and sometimes opposing, effects on cellular growth and inflammatory signalling pathways, consistent with our new appreciation of the regulation of KapB translation by *kaposin* NS retention. This work also provides a valuable roadmap for how to disentangle the complex myriad of functions elicited by bifunctional transcripts that are encoded by both cellular and viral genes.

## RESULTS

### Characterization of an overexpression model to disentangle kaposin RNA and KapB functions

We recently characterized our kaposin constructs in HEK-293T transfection [38], and when we immunoblotted for αKap DRs, several unknown protein products were detected with ΔStarts. In transfection protocols, cells receive an unbalanced dose of genetic material that could cause aberrant protein products to be produced [43, 44]. While this could provide hints into intriguing biological phenomena, we instead sought to develop a reductionist model that better mimics *kaposin* transcription and translation in infection. For this, we employed lentivirus transduction to overexpress our kaposin constructs which achieves stable albeit random integration of the gene of interest into the host genome that is passed onto cell progeny [45].

To characterize our overexpression model, we transduced A549 cells with recombinant lentiviruses expressing kaposin constructs alongside an empty vector (EV) control and immunostained for NS-resident protein, SRRM2, and Kap B/C using an anti-Kap DRs polyclonal antibody [38] followed by RNA-FISH for *kaposin* RNA as in [38] (**Fig. 1C**). Irregularly shaped nuclear speckles were observed in EV control cells without any detectable Kap proteins or *kaposin* RNA. Both WT and ΔStarts overexpression resulted in the formation of *kaposin* foci that strongly colocalized with SRRM2, as expected [38]. A small amount of KapB/C protein was detected in the nucleus after WT overexpression but not in cells expressing ΔStarts. Since we have not isolated for a single-cell clone, we observed heterogeneity in the cell population where cells display different expression levels of *kaposin* RNA (WT, ΔStarts) or Kap proteins (WT only). By contrast, lentiviral transduction of AltB constructs showed that this recoded transcript failed to form *kaposin*-NSs because AltB transcripts lack the GC-rich DRs required for *kaposin*-NS formation. However, AltB did result in the most Kap B protein expression [38]. We noted the same heterogeneity in AltB overexpressing cells for Kap protein detection. In parallel, we harvested protein lysates from A549 cells transduced with kaposin constructs then immunoblotted using αKap DRs (**Fig. 1D**). This antibody recognizes KapB and KapC, since they contain Kap DRs, and not KapA. We detected a strong band in AltB overexpression at ∼45 kDa which matches the predicted molecular weight of KapB. Weak yet detectable amounts of KapB was observed in WT, which is more prominent on an overexposed blot. The faint band above KapB in WT lysates likely represents KapC. These results are consistent with the expectation that most of the WT transcript is retained in NSs, with little RNA exported to the cytoplasm for translation. Neither KapB nor KapC was detected after ΔStarts overexpression.

To confirm that the limited or lack of Kap protein expression in WT and ΔStarts was not due to differences in *kaposin* transcript abundance, we harvested total RNA from parallel wells from transduced cells and performed RT-qPCR using a TaqMan assay specific for ORFK12/KapA sequences which remain unchanged in all kaposin constructs. We observed that overexpression of all kaposin constructs resulted in a change in total cell yield relative to EV; specifically, WT and ΔStarts increased cell yield whereas AltB decreased cell yield relative to EV and WT (**Fig. 1E**). Because of this difference, we estimated the total number of *kaposin* transcripts detected across constructs then normalized it on a per cell basis (**Fig. 1F**). While no *kaposin* transcripts were detected in control cells, *kaposin* was detected in cells transduced with WT, ΔStarts, and AltB, without any significant difference in transcript number between transductions (**Fig. 1F**). These data are consistent with the previously observed NS retention of *kaposin* during WT and ΔStarts overexpression, limiting translation of these RNAs. By contrast, AltB *kaposin* is not retained in NSs, and is translated more abundantly. Overall, we have created a lentiviral transduction model that uncouples the overexpression of *kaposin* RNA from KapB protein that more closely mimics the infection state. This system allows us to re-assess whether phenotypes previously attributed to KapB protein are instead a function of *kaposin* RNA.

### KapB protein and not kaposin RNA induce stress fibre formation and changes in cell size

Previous work found that KapB overexpression in primary endothelial cells induced actin remodelling and stress fibre formation, eliciting changes in cell shape to an elongated, spindled morphology [25, 28]. To assess if kaposin RNA or KapB protein was responsible for this effect, we transduced A549 cells with our panel of lentiviruses and stained for actin stress fibres using phalloidin which binds filamentous actin alongside eGFP as a protein overexpression control. Actin remodelling and stress fiber formation was not obvious in this lung epithelial cell line [46]. Next, we transduced the endothelial hybrid cell line EA.hy926 [47, 48] with kaposin constructs including an AltB construct where KapB is FLAG-tagged (AltB-FLAG); for this construct, KapB expression was detected with αFLAG immunofluorescent staining. EA.hy926 cells are commonly used to monitor cytoskeleton remodeling and actin SF formation [49, 50]. We visually observed that overexpression with AltB-FLAG, but not ΔStarts, induced actin stress fibres and changed cell shape (**Fig. 2A**). To quantify changes in actin, we used Fiji to measure changes in actin intensity from a line drawn across cells (**Fig. 2B**). We found no obvious changes in the actin intensity profile when comparing EV, high *kaposin* expressing ΔStarts cells (based on cells displaying visible *kaposin* NSs), and high eGFP expressing cells (based on cells with abundant eGFP detection). AltB-FLAG overexpressing cells that stained positive for FLAG (high KapB expressors) displayed a drastic increase in actin intensity amplitude and frequency (**Fig. 2B**). This data suggests that KapB protein, and not *kaposin* RNA, mediates actin rearrangements and induces stress fibre formation in cells.

**Figure 2.**
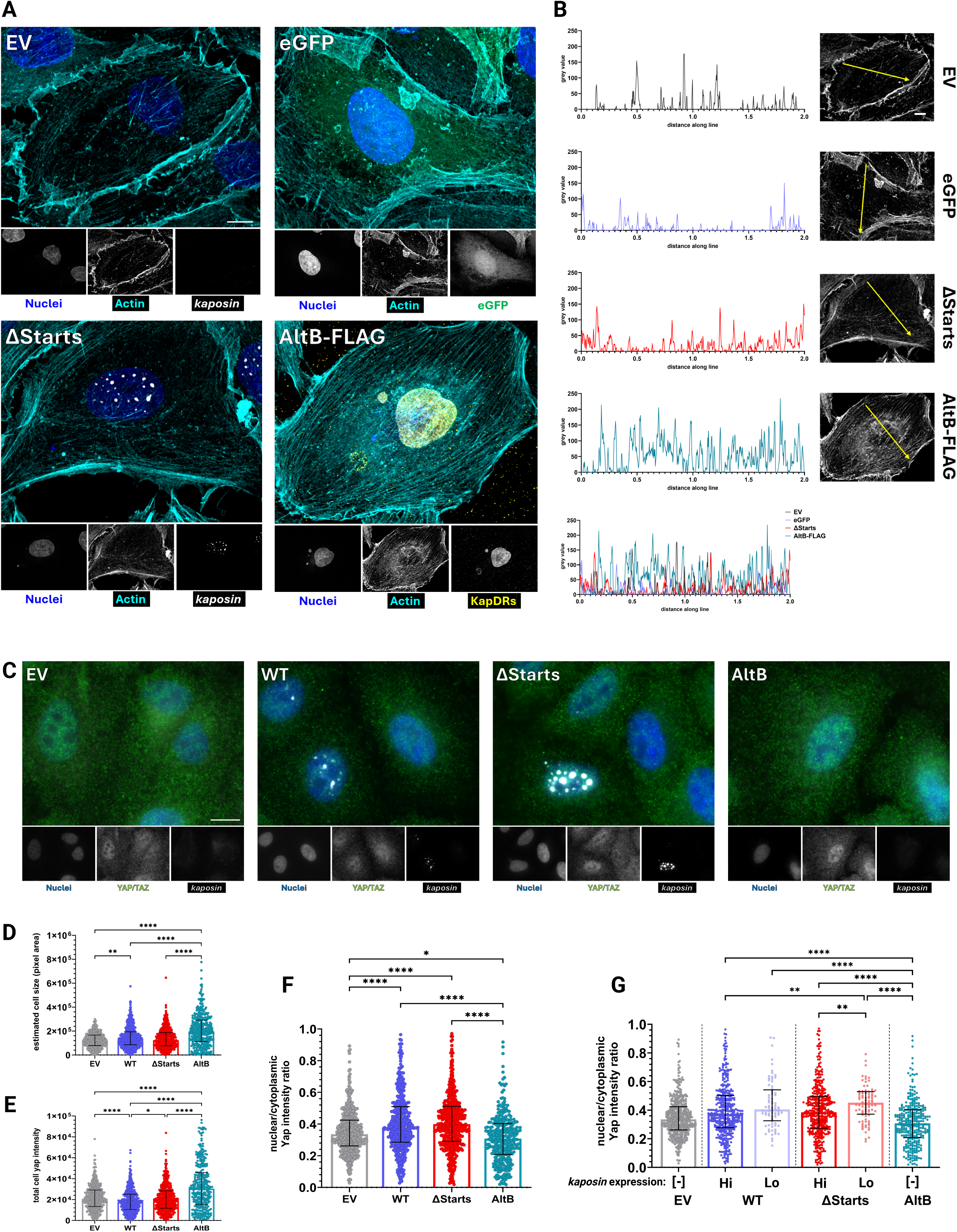
Kaposin RNA induces and KapB protein restricts YAP/TAZ translocation. **(A)** EA.hy926 cells were transduced with ΔStarts and AltB-FLAG alongside EV and eGFP controls. Post-selection, cells were fixed and either immunostained for FLAG (AltB-FLAG only to detect KapB; false-coloured yellow; AlexaFluor-488) or RNA-FISH for *kaposin* RNA (EV and ΔStarts only; false-coloured white; fluorescein-495). All conditions include a Hoechst stain for nuclei (false-coloured blue) and phalloidin-647 to visualize F-actin (false-coloured cyan). Photomicrographs were taken using a Zeiss LSM 880 Airyscan confocal microscope and presented as maximum intensity projections. One representative image set of three independent replicates (*n* = 3) is shown, and independent channels are shown in grayscale. Scale bar = 10 μm. **(B)** Grayscale actin channel images from Figure 2A were analyzed using ImageJ to plot actin intensity profiles along the indicated lines shown in yellow. One representative actin intensity profile set of three independent replicates (*n* = 3) is shown. **(C)** A549 cells were transduced with WT, ΔStarts, and AltB alongside an EV control; post-selection, cells were fixed and immunostained for YAP/TAZ (false-coloured green; AlexaFluor-647) followed by RNA-FISH for *kaposin* (false-coloured white; fluorescein-495) and Hoechst stain for nuclei (false-coloured blue). Photomicrographs were taken using a Zeiss AxioObserver Z1 microscope. One representative image set of three independent replicates (*n* = 3) is shown, and independent channels are shown in grayscale. Scale bar = 10 μm. **(D-F)** Grayscale YAP channel images from Figure 2C were analyzed using CellProfiler to approximate cell borders based on local changes in YAP/TAZ staining intensity. These borders were used to approximate cell size expressed as pixel area, total YAP/TAZ intensity expressed as pixel intensity, and nuclear/cytoplasmic YAP/TAZ ratios. Each dot represents a single cell, and at least 107 cells were measured per replicate. Statistical analysis was conducted using a Kruskal-Wallis H test with Dunn’s post-hoc (*n* = 3; median with interquartile range; * *p* < 0.05, ** *p* < 0.01, **** *p* < 0.0001). **(G)** CellProfiler was used to stratify nuclear/cytoplasmic YAP/TAZ ratios from Figure 2F based on cells displaying high (Hi) or low (Lo) *kaposin* RNA expression in WT and ΔStarts overexpressing cells. Cells overexpressing EV and AltB are denoted negative [–] for GC-rich, repetitive *kaposin*. Statistical analysis was conducted using a Kruskal-Wallis H test with Dunn’s post-hoc (**n** = 3; median with interquartile range; ** *p* < 0.01, **** *p* < 0.0001).

We previously found that KapB-mediated actin rearrangements and stress fibre formation were required for the activation of the pro-growth transcriptional co-activator yes-associated protein (YAP) [28]. A549 cells were transduced with kaposin constructs then immunostained using an antibody that recognizes total YAP and TAZ followed by RNA-FISH for *kaposin* (**Fig. 2C**). We again observed that AltB appeared to alter cell size, and so YAP/TAZ staining was used to approximate cell borders based on local changes in YAP/TAZ intensity (**Fig. 2D**). This serves only as a proxy to estimate cell size since we did not conduct a stain to define cell borders. We found that cells overexpressing AltB and producing abundant KapB had a significantly greater estimated cell size compared to all other constructs. Cells overexpressing the WT construct translate a small amount of KapB and displayed a greater estimated cell size compared to EV but reduced cell size compared to AltB. There was no change in estimated cell size between EV and ΔStarts, suggesting that only KapB and not *kaposin* influences cell size.

### Kaposin RNA induces YAP/TAZ nuclear translocation while KapB protein restricts it

Using the same samples stained for total YAP, we also measured total cell YAP/TAZ intensity (**Fig. 2E**). Overexpression with AltB resulted in significantly greater YAP/TAZ intensity compared to all other constructs, whereas WT resulted in a significant reduction in YAP/TAZ intensity compared to all other constructs. This data suggests that KapB protein alone either induces YAP/TAZ expression or reduces their degradation which is partially reverted in the presence of *kaposin* RNA. Though measuring total YAP/TAZ intensity can help to infer their regulation, we are unable to draw conclusions on YAP/TAZ activation status from this measurement. To do this, we separated intensity measurements based on nuclear or cytoplasmic compartments then took the ratio of YAP/TAZ intensity between these (i.e., nuclear/cytoplasmic; **Fig. 2F**). Since nuclear YAP/TAZ intensity is in the numerator, an increase in this ratio would suggest the nuclear translocation of YAP/TAZ, thereby inferring their activation. We found that WT and ΔStarts overexpression resulted in a significantly greater nuclear/cytoplasmic YAP/TAZ intensity ratio relative to EV and AltB, suggesting they have relatively increased YAP/TAZ nuclear translocation. AltB overexpression instead resulted in a significant reduction in this ratio relative to all other constructs, suggesting that KapB is restricting YAP/TAZ nuclear translocation.

To address the heterogeneity of this phenotype, we modified our imaging analysis to stratify WT and ΔStarts cells based on high *kaposin* expression (i.e., cells with highly visible *kaposin* foci) or low *kaposin* expression (i.e., cells with low or no visible *kaposin* foci). Since EV and AltB do not produce GC-rich *kaposin* RNA, we denoted them to have negative (–) *kaposin* expression. When the results were re-analyzed to apply this stratification (**Fig. 2G**), we found that WT and ΔStarts cells with high *kaposin* expression had the greatest significant increase in nuclear/cytoplasmic YAP/TAZ intensity ratio relative to all other comparisons. Furthermore, there was no significant change in this ratio in WT and ΔStarts cells with low *kaposin* expression compared to EV. Cells overexpressing AltB had the greatest reduction in this ratio relative to all other comparisons. These results support the notion that KapB protein and *kaposin* RNA may differentially mediate YAP/TAZ signalling: *kaposin* RNA induces YAP/TAZ translocation while KapB protein restricts it but increases total YAP abundance.

### Kaposin RNA and KapB protein differentially modulate YAP biochemical regulation and cellular metabolism

Another method used to infer YAP/TAZ activation is by assessing YAP/P-YAP and TAZ/P-TAZ levels via immunoblot. Since only unphosphorylated YAP/TAZ can translocate into the nucleus to activate YAP/TAZ-target genes, increasing YAP/P-YAP or TAZ/P-TAZ ratios suggests their activation [51, 52]. To monitor this, protein lysates from A549 transduced with kaposin constructs were harvested then used to immunoblot for YAP and P-YAP (**Fig. 3A**) or TAZ and P-TAZ (**Fig. 3C**) followed by densitometry analysis. There was no significant changes in total YAP or P-YAP levels across kaposin constructs relative to EV; however, AltB overexpression caused a significant increase in total YAP/P-YAP ratio relative to EV (**Fig. 3B**), suggesting that KapB induces YAP activation. This also corresponds with the microscopy data showing that KapB increases total YAP levels (**Fig. 2E**). Meanwhile, all kaposin constructs resulted in a significant reduction in both TAZ and P-TAZ levels with no significant change in TAZ/P-TAZ ratio relative to EV (**Fig. 3D**). This would suggest that overexpression of all kaposin constructs either directly or indirectly induces the degradation of TAZ, further suggesting that YAP and not TAZ is integral for KapB effects.

**Figure 3.**
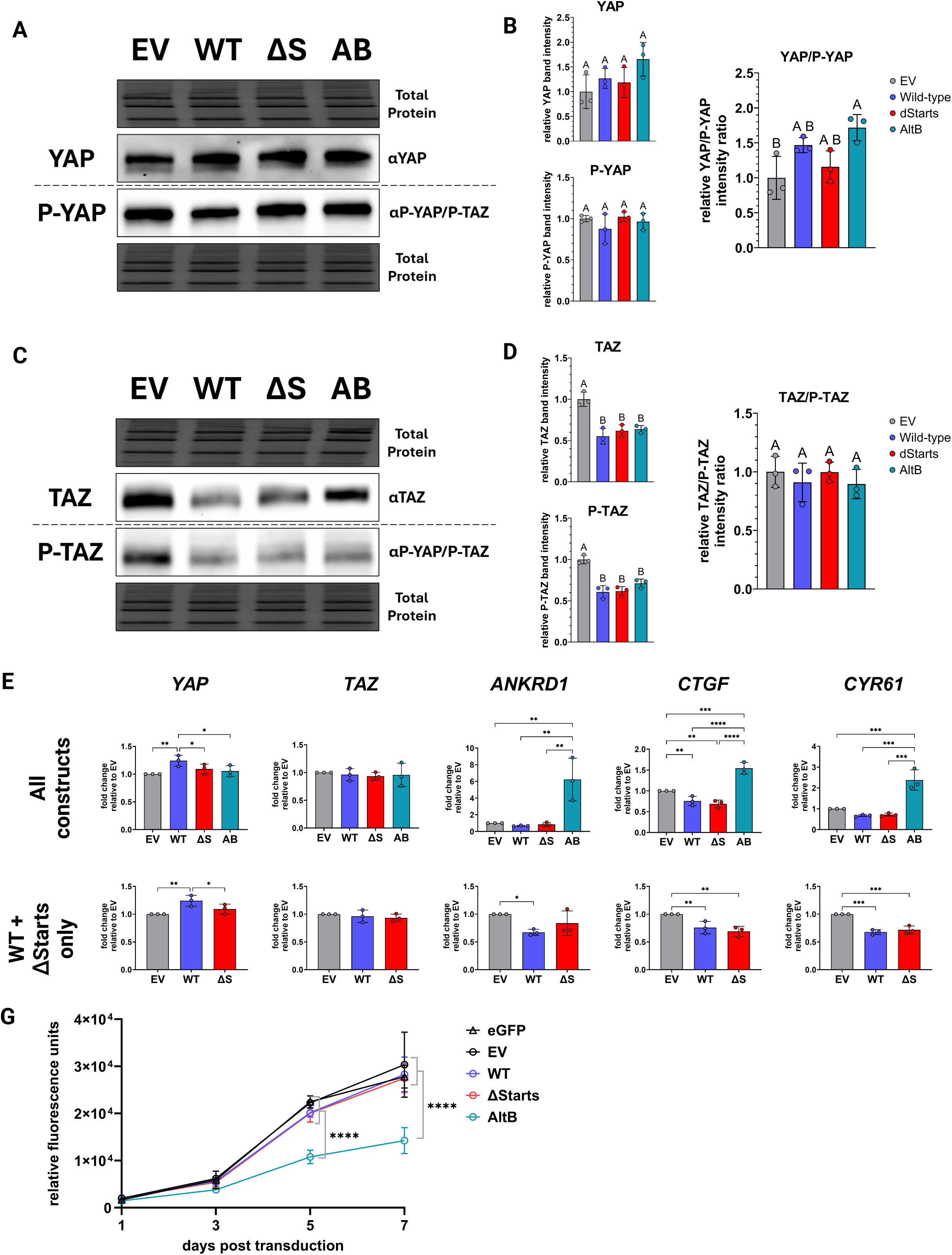
KapB protein, and not kaposin RNA, reduces cellular metabolic activity. **(A-D)** A549 cells were transduced with WT, ΔStarts, and AltB alongside an EV control; post-selection, protein lysates were harvested and used to immunoblot for antibodies specific for YAP, TAZ, and P-YAP and P-TAZ. One representative blot of three independent replicates (*n* = 3) is shown. Densitometry analysis was performed to quantify YAP, P-YAP, TAZ, and P-TAZ band intensity, normalized to total protein loading control, then intensity values used to calculate YAP/P-YAP and TAZ/P-TAZ ratios. All data are expressed as a fold change relative to EV. Statistical analysis was conducted using an ordinary one-way ANOVA with a Tukey’s post-hoc (*n* = 3; mean ± SEM; compact letter display where different lettering indicates *p* < 0.05). **(E)** RT-qPCR was performed on cDNA generated from A549 cells transduced with WT, ΔStarts, and AltB alongside an EV control. Relative transcript levels of *YAP*, *TAZ*, *CTGF*, *CYR61*, and *ANKRD1* were determined via ΔΔCq analysis using *HPRT* as a reference gene. Data are expressed as a fold change relative to EV. Statistical analysis was conducted using a repeated-measures one-way ANOVA with an uncorrected Fisher’s least significant difference post-hoc (*n* = 3; mean ± SEM; * *p* < 0.05; ** *p* < 0.01; *** *p* < 0.001; **** *p* < 0.0001). **(F)** A549 cells were transduced with WT, ΔStarts, and AltB alongside an EV and eGFP control. Cell metabolic activity was assessed at 1, 3, 5, and 7 days post-transduction using a resazurin assay (*n* = 3; mean ± SEM). Statistical analysis was conducted using a two-way repeated-measures ANOVA with a Bonferonni correction (**** *p* < 0.0001).

As another measure of YAP activation, we quantified the steady-state levels of *YAP* and *TAZ* transcripts, and the transcripts of YAP-target genes, *ANKRD1*, *CTGF*, and *CYR61*. There was no appreciable change in *YAP* and *TAZ* transcript levels across kaposin constructs (**Fig. 3E**), suggesting that changes in YAP and TAZ protein levels (**Fig. 3A-D**) occur post-transcriptionally. Overexpression with AltB resulted in an overall significant increase in YAP-target gene transcript abundance, whereas WT and ΔStarts resulted in their significant reduction (**Fig. 3E**). This data suggest that KapB is activating pro-growth signalling that is restricted in the presence of *kaposin* NSs.

Collectively, KapB appears to be activate YAP signalling, as seen by increase in total YAP intensity, increase in total YAP/P-YAP ratio, and increase in YAP-target genes. However, we noted earlier that AltB results in a strongly significant reduction in total cell yield compared to other constructs (**Fig. 1E**). Previous reports have suggested that overexpression of KapB induces cytotoxic effects [10, 20, 22], raising the possibility that reduced cell yield is due to impaired cell growth or viability. Cellular metabolic activity was assessed using a resazurin assay (**Fig. 3G**). In the presence of cells, resazurin is metabolically reduced to resorufin that is detected by fluorometry, which is used an indirect measure of viable cells [53–55]. By 5 days post-transduction, AltB overexpression resulted in a significant reduction in cellular metabolism relative to all other constructs, including an eGFP control for protein overexpression. This reduction was sustained by 7 days post transduction. These data suggest that KapB causes a reduction in metabolically active cells, hinting that KapB and not *kaposin* might induce cell growth arrest or cytotoxic effects.

### Kaposin RNA induces cell autonomous p-body disassembly

We previously found that YAP was required for KapB/*kaposin* induced processing (p-)body disassembly [28]. P-bodies are ribonucleoprotein (RNP) granules constitutively present in the cytoplasm that regulate the stability, turnover or decay of AU-rich element (ARE-)mRNAs [56, 57]. ARE-mRNAs encode for pro-inflammatory, pro-growth, and pro-angiogenic biomolecules, whose transcripts become available for translation upon p-body disassembly. Armed with the knowledge that *kaposin* RNA and KapB protein differentially regulate YAP signalling, we used our overexpression model to monitor the contribution of *kaposin* RNA and KapB protein to p-body disassembly. A549 (**Fig. 4A,B**) and EA.hy926 (**Fig. 4C,D**) were transduced with kaposin constructs then immunostained for p-body resident DEAD-box helicase 6 (DDX6) followed by RNA-FISH for *kaposin* (**Fig. 4A,C**). All kaposin constructs caused a significant reduction in DDX6 foci/cell in the total cell population relative to EV, with WT and ΔStarts overexpression resulting in the greatest reduction in both cell types (**Fig. 4B, D**). These data suggest that *kaposin* RNA may elicit p-body disassembly. The presence of visible *kaposin* foci negatively correlated with p-bodies in WT and ΔStarts overexpressing cells (**Fig. 4A, C**). To confirm this, data from Figure 4B and 4D were stratified based on high and low *kaposin* RNA expression as in Figure 2G (**Fig. 4C**). All conditions resulted in a significant reduction in DDX6 foci/cell relative to EV. However, cells displaying high *kaposin* expression in WT and ΔStarts had significant DDX6 foci/cell reduction compared to AltB overexpression or cells with low *kaposin* expression (**Fig. 4B,D**). Notably in endothelial EA.hy926 cells (**Fig. 4D**), low *kaposin* expressing WT and ΔStarts cells show no change in DDX6 foci/cell compared to AltB. These results suggest that overexpression of *kaposin* RNA elicits p-body disassembly in a cell autonomous manner and is not cell-type specific (epithelial A549, or endothelial EA.hy926).

**Figure 4.**
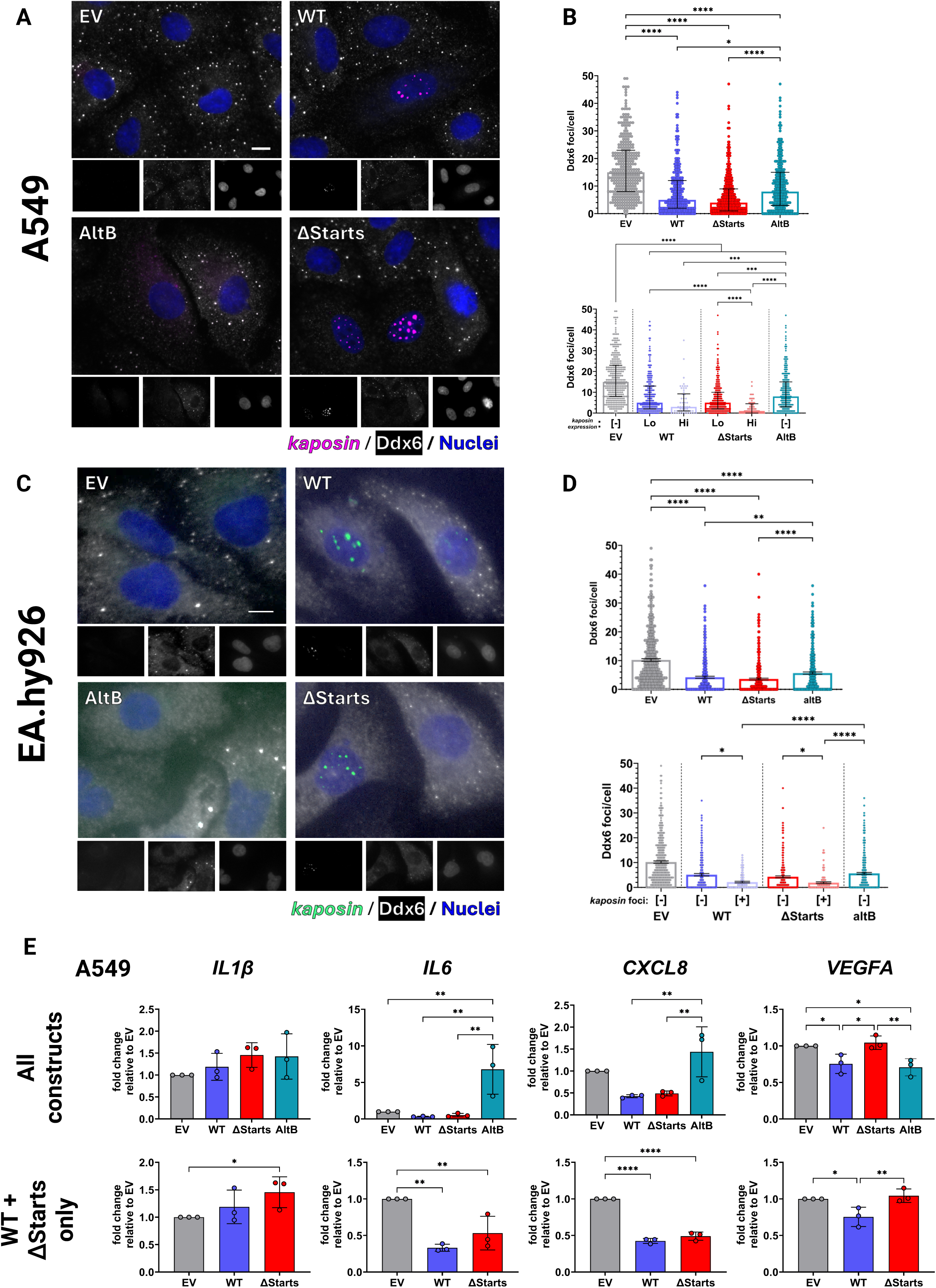
Kaposin RNA induces cell autonomous p-body disassembly. **(A, B)** A549 and **(C, D)** EA.hy926 cells were transduced with WT, ΔStarts, and AltB alongside an EV control; post-selection, cells were fixed and immunostained for DDX6 (false-coloured white; AlexaFluor-647) followed by RNA-FISH for *kaposin* (false-coloured magenta in A549 cells and false-coloured green in EA.hy926 cells; fluorescein-495) and Hoechst stain for nuclei (false-coloured blue). Photomicrographs were taken using a Zeiss AxioObserver Z1 microscope. One representative image set of three (A549, *n* = 3) or four (EA.hy926, *n* = 4) independent replicates (*n* = 4) is shown, and independent channels are shown in grayscale. Scale bar = 10 μm. Grayscale DDX6 channel images were analyzed to enumerate the number of DDX6 foci on a per cell basis in the total cell population and with cells stratified based on *kaposin* expression level as in **Fig. 2G**. Each dot represents a single cell, and at least 102 cells were measured per replicate. Statistical analysis was conducted using a Kruskal-Wallis H test with Dunn’s post-hoc (*n* = 4; median with interquartile range; * *p* < 0.05, ** *p* < 0.01, **** *p* < 0.0001). *(E)* RT-qPCR was performed on cDNA generated from A549 cells transduced with WT, ΔStarts, and AltB alongside an EV control. Relative transcript levels of *IL-1B*, *IL-6*, *CXCL-8* and *VEGFA* was determined via ΔΔCq analysis using *18S rRNA* as a reference gene. Data are expressed as a fold change relative to EV. Statistical analysis was conducted using a repeated-measures one-way ANOVA with an uncorrected Fisher’s least significant difference post-hoc (*n* = 3; mean ± SEM; * *p* < 0.05; ** *p* < 0.01; **** *p* < 0.0001).

To assess the link between p-body disassembly and inflammation, we measured the steady-state transcript levels of p-body regulated cytokine ARE-mRNAs, *IL-1β, IL-6, CXCL-8* and *VEGFA*; these transcripts were also reported to be YAP target genes [58]. There was no appreciable change in *IL-1β* or *VEGFA* transcript levels across kaposin constructs (**Fig. 4E**). AltB overexpression resulted in a significant increase in *IL-6* and *CXCL-8* transcript abundance, whereas WT and ΔStarts resulted in a significant reduction relative to EV (*Fig. 4E*). This data suggest that KapB is activating pro-inflammatory signalling that is restricted in the presence of *kaposin* NSs.

### KapB antagonizes p-bodies using autocrine and paracrine signalling

Early studies probing the function of KapB found that KapB utilizes DR2 amino acid repeats to bind to and activate mitogen-activated protein kinase-activated protein kinase 2 (MAPKAPK2 or MK2), leading to the stabilization or reduced degradation of ARE-mRNAs [20, 21]. This resulted in a potent inflammatory response and release of cytokines in cell supernatants such as GM-CSF and IL-6 which may activate autocrine or paracrine signalling pathways [20]. Previous studies have linked the rearrangement of p-bodies to the stabilization of the ARE-mRNAs that encode these cytokines [59, 60]. To test this phenomenon, we first transduced EA.hy926 cells with kaposin constructs, alongside a no transduction control and an eGFP protein overexpression control, then harvested cell-conditioned media (CM). KS lesions are composed of ECs; therefore, we used EA.hy926 for this experiment as these cells retain endothelial inflammatory functions and respond to paracrine signalling [61, 62]. Naïve, non-transduced EA.hy926 cells were sensitized by starvation for 2 h then treated with CM for 2 h followed by immunostaining for DDX6 and YAP/TAZ (**Fig. 5A**).

**Figure 5.**
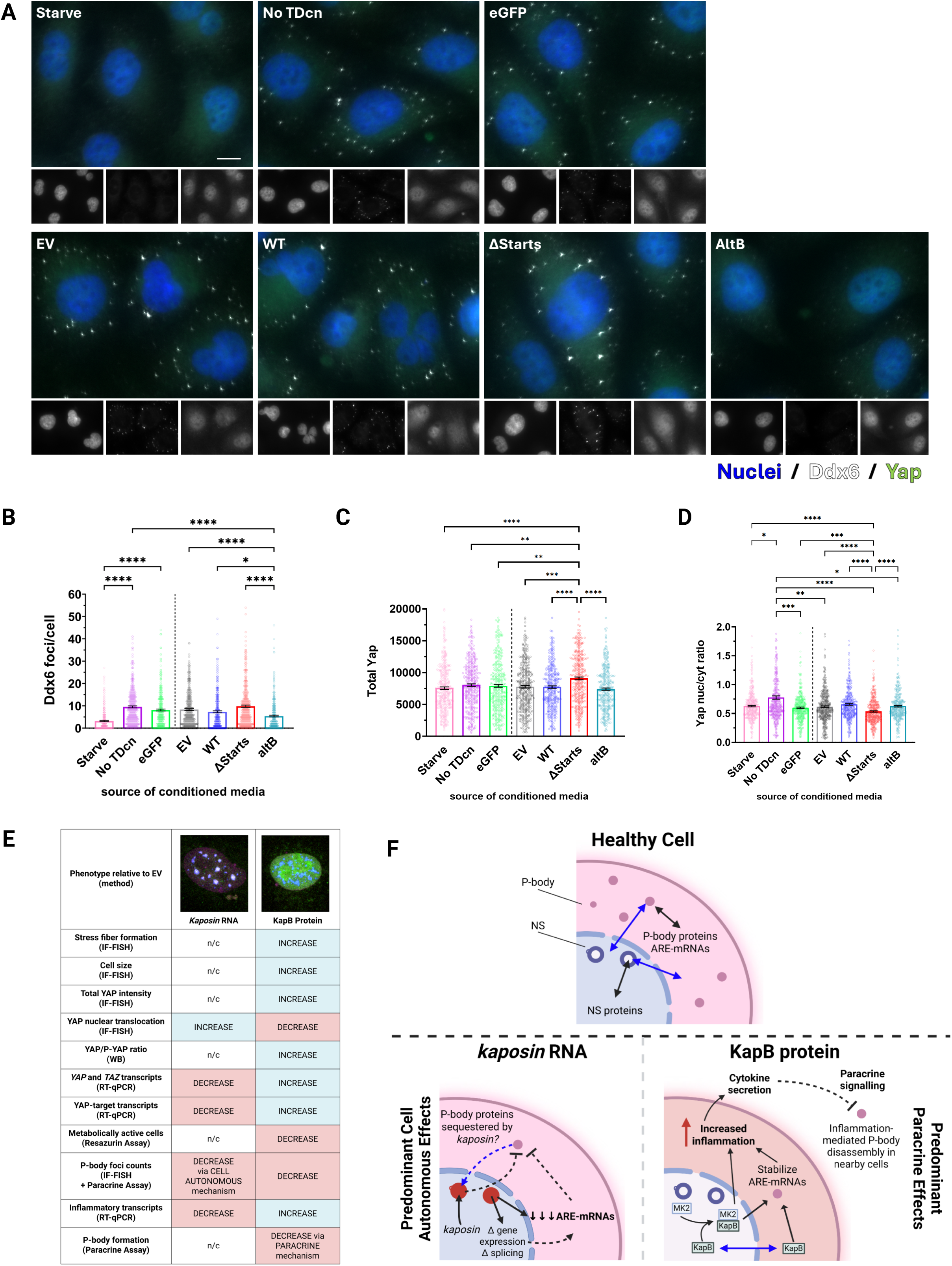
KapB-induced secretory factors antagonize p-body restoration. **(A-D)** EA.hy926 cells were either not transduced or transduced with WT, ΔStarts, and AltB alongside EV and eGFP controls. Post-selection, cells were starved for 48 h to generate conditioned media (CM). Naïve EA.hy926 cells were sensitized by starvation for 2 h, treated with CM for 2 h, then cells were fixed and immunostained using antibodies specific for YAP/TAZ (false-coloured green; AlexaFluor-488) and DDX6 (false-coloured white; AlexaFluor-647), and Hoechst stain for nuclei (false-coloured blue). CellProfiler was used to enumerate the number DDX6 foci/cell as in **Fig. 4B**, total YAP/TAZ intensity as in **Fig. 2E**, and nuclear/cytoplasmic YAP/TAZ ratio as in **Fig. 2F**. Each dot represents a single cell, and at least 115 cells were measured per replicate. Statistical analysis was conducted using a Kruskal-Wallis H test with Dunn’s post-hoc (*n* = 3; median with interquartile range; * *p* < 0.05, ** *p* < 0.01, *** *p* < 0.001, **** *p* < 0.0001). **(E)** Summary of *kaposin* RNA and KapB phenotypes assessed in this study. Light blue coloured panels indicate increases relative to EV, and light red coloured panels indicate decreases relative to EV. **(F)** *Kaposin* RNA and KapB protein use different mechanisms to induce p-body disassembly. Solid arrows indicate known phenotypes, and dashed arrows indicate proposed mechanisms for unknown phenotypes. (i) In healthy cells, p-bodies and NSs are in constant exchange with their surrounding environment (black arrows). Select p-body and NS resident proteins are capable of shuttling between the cytoplasm and nucleus (blue arrows). (ii) *Kaposin* RNA acts as a scaffold to form nuclear *kaposin* foci containing NS machinery and induces cell autonomous p-body disassembly through an unknown mechanism. One possibility is through the reduction in steady-state levels of ARE-mRNAs. Another possibility is that the formation of *kaposin* foci sequesters p-body resident proteins away from the cytoplasm, enabling their disassembly. (iii) KapB-mediated inflammation, via MK2 activation and p-body disassembly, may induce the release secretory molecules, like inflammatory cytokines, that antagonize p-bodies in neighbouring cells through paracrine signalling.

Starvation causes a reduction in DDX6 foci/cell which was restored when treated with CM from non-transduced cells or from cells overexpressing eGFP (**Fig. 5B**). CM from cells overexpressing EV, WT, and ΔStarts resulted in complete restoration of DDX6 foci/cell i.e., no change relative to the no transduction and eGFP controls. However, CM collected from cells overexpressing AltB resulted in only a partial restoration in p-bodies, as seen by a significant increase in DDX6 foci/cell relative to starvation but a significant reduction relative to all other conditions. These data suggest that KapB induces the release of secretory factors that inhibit the formation of p-bodies. Surprisingly, CM from ΔStarts overexpression resulted in a significant increase in YAP/TAZ intensity but a decrease in nuclear/cytoplasmic YAP/TAZ ratio compared to all other conditions (**Fig. 5C,D**), suggesting that **kaposin** might induce the release of secretory factors that restrict YAP/TAZ nuclear translocation in bystander cells. Taken together, these results demonstrate that KapB activates a paracrine signalling axis inhibitory to p-bodies, likely due to the activation of cellular growth and inflammatory signalling in these cells.

## DISCUSSION

The KSHV kaposin gene has historically been viewed as a protein-coding mRNA, with investigations of Kap proteins heavily biased towards the abundant KapB protein [10]. Our discovery that *kaposin* RNA is largely retained in the nucleus where it has a non-coding function that supersedes that of its protein products [38] reframes all prior work that was performed using overexpression models that produce both KapB and *kaposin*. In the current study, we developed a lentivirus overexpression system (*Fig. 1B-F*) to disentangle the contributions of *kaposin* RNA and KapB protein to cellular phenotypes previously attributed to KapB overexpression [20, 25, 28]. We now show that *kaposin* RNA and KapB protein mediate distinct effects to differentially modulate cellular growth and inflammatory signalling (*Fig. 5E*). We propose a that *kaposin* RNA induces cell autonomous p-body disassembly with a concomitant reduction in the steady-state levels of growth and inflammatory transcripts (**Fig. 5F**) while KapB activates growth and inflammatory signalling to antagonizes p-bodies in a paracrine manner (**Fig. 5F**) despite also reducing cellular metabolic activity.

### A reductionist overexpression system mimics kaposin and Kap protein expression during different KSHV infection phases

The default replication phase after KSHV primary infection is viral latency, where the viral episome is largely transcriptionally silenced save for the latency locus, no virus is produced, and host cells survive [63–65]. In response to environmental or cellular stresses, such as hypoxia and oxidative stress [66, 67], the virus can reactivate, a phase characterized by complete viral gene expression, virion production, virion release and cell death [63, 65]. *Kaposin* has traditionally been considered a latent gene based on abundant reports of *kaposin* transcript expression during latent infection or in latently infected cells in KS lesions [7–15, 38]. Thus, when *kaposin* was discovered to be a polycistronic, protein-coding mRNA encoding for KapA, B and C [10], Kap proteins were presumed to be expressed as latent proteins. Recent work from our lab has changed this paradigm. Using RNA fluorescent *in situ* hybridization (FISH), we discovered that *kaposin* is largely retained in NSs during latent infection [38]. The lack of kaposin mRNA export during this infection phase means that Kap proteins are either not expressed or minimally expressed during latency. By contrast, after lytic reactivation and enhanced *kaposin* transcription, RNA FISH revealed that although *kaposin* is found in NSs, *kaposin* can also be visualized in the cytoplasm, consistent with the detection of abundant KapB production during late lytic infection [38]. Taken together, these data suggest that although *kaposin* is a latent transcript, KapB (and by inference, KapA and KapC) are lytic proteins and are not or minimally expressed during latency. In the current study, we have endeavored to mimic these stages of KSHV infection using a reductionist system (**Fig. 1C,D**). With ΔStarts, a construct that encodes the authentic RNA molecule but fails to translate Kap A, B, C proteins, we model KSHV latent infection. With AltB, a construct that encodes the authentic amino acid sequence of KapB from an RNA that lacks the DRs nuclear-retention signal, we mimic the extreme state of abundant KapB expression that occurs in late lytic reactivation. As well as acting as simplified models for *kaposin* expression in different KSHV phases, with these two constructs we aimed to uncouple functions elicited by KapB from those elicited by *kaposin*. We validated that each overexpression construct behaved as expected in terms of NS retention and protein translation and ruled out the possibility that any difference in protein expression was due to differences in transcript levels across constructs (**Fig. 1E,F**). Here, we have created a reductionist model to accurately uncouple **kaposin** function from that of KapB while also providing a simplified system to mimic *kaposin* versus KapB expression during KSHV latent and lytic replication phases.

### Kaposin and KapB mediate opposing actin remodelling and YAP regulation phenotypes

Yes-associated protein (YAP) is a mechanosensitive transcriptional co-activator and central player of the Hippo signalling pathway, which controls cellular proliferation and growth (reviewed in [51, 68]). YAP must be translocated into the nucleus and associate with nuclear co-transcriptional co-activators to drive pro-proliferative and pro-growth gene expression. This is biochemically regulated by upstream Hippo kinases; YAP can translocate into the nucleus only in its unphosphorylated form [51, 68]. In response to environmental signals where cell growth and proliferation are not favoured, like serum starvation [69], YAP is phosphorylated (P-YAP) by Hippo kinases and either retained in the cytoplasm or targeted for degradation [70, 71]. This is also true for the YAP paralog transcriptional co-activator with PDZ-binding motif (TAZ), also known as WW domain-containing transcription regulator protein 1 (WWTR1) [72, 73]. Overexpression with all kaposin constructs led to differential biochemical regulation of YAP and TAZ, where total YAP or P-YAP levels were unchanged or increased (**Fig. 3A,B**) and total TAZ or P-TAZ levels decreased (**Fig. 3C,D**). Since there was no change in *YAP* or *TAZ* transcript levels across constructs (**Fig. 3E**), we reason that both *kaposin* and KapB may be mediating the protein degradation of TAZ/P-TAZ. This suggests that YAP, and not TAZ, is important in kaposin overexpression.

Though the biochemical regulation of YAP and TAZ is well defined, there is growing appreciation that these biomolecules are mechanically regulated in a manner distinct from traditional Hippo signalling. Cells exposed to mechanical cues, such as shear stress and extracellular matrix stiffness, induces actin SF formation and changes in cell size and tension [74–76]. These mechanical forces pull on the linker of nucleoskeleton and cytoskeleton (LINC) complexes, using actinomyosin contractility, to open nuclear pores and induce YAP nuclear translocation independent of upstream Hippo regulators [74–78]. In prior studies using KapB overexpression in primary ECs, KapB-mediated actin rearrangements, SF formation, and actinomyosin contractility were required for the activation of YAP [28]. KapB caused and overall increase in total YAP levels as detected by immunoblot but resulted in no change in nuclear YAP translocation as assessed by microscopy [28]. When YAP was knocked down, KapB was no longer able to induce SF in cells [28], demonstrating that YAP is required for KapB-mediated SF formation. We now show that KapB induces cell-specific SF formation (**Fig. 2A,B**) that corresponds with increased total YAP levels (**Fig. 3A,B**), along with a potent increase in the transcription of YAP-target genes (**Fig. 3E, 4E**). These data confirm that KapB activates YAP signaling, consistent with our previous observations [28]. To confirm that KapB-mediated YAP activation requires SF formation, future studies will test whether AltB activates YAP-target gene transcription when actin contractility is disrupted using blebbistatin [28, 79].

The LINC complex consists of nesprin scaffolding proteins found in the outer nuclear membrane that connects the actin cytoskeleton to the nuclear lamina through SUN proteins [78, 80]. In response to mechanical stimuli, the LINC complex gets pulled and causes nuclear flattening to increase nuclear pore opening and YAP translocation [76, 81–83]. When the LINC complex was disrupted using a dominant negative nesprin construct, actinomyosin contractility from mechanical stimuli was no longer able to induce YAP nuclear translocation [81]. We showed that KapB did not increase the ratio of nuclear to cytoplasmic YAP (**Fig. 2F,G**), despite its ability to induce SF formation. This is unexpected, as SF formation mediates nuclear flattening and YAP translocation through LINC tension. These findings are inconsistent with our previous observations [28]; however, this former study used a construct that produced both *kaposin* and KapB, making it possible that these inconsistencies are due to variable expression level of KapB. Conversely, we showed that *kaposin* increase nuclear YAP translocation in a cell autonomous manner (**Fig. 2G**) despite its inability to induce SF formation (**Fig. 2A,B**). Taken together, KapB and *kaposin* appear to regulate nuclear YAP translocation in opposing ways. Future studies will explore whether *kaposin* or KapB require the LINC complex to mediate differences in nuclear YAP translocation.

### Dual regulation of p-bodies by a bifunctional RNA and its protein product

P-bodies are constitutively present, cytoplasmic ribonucleoprotein (RNP) granules that regulate mRNA turnover, possessing enzymes with activities for 5’-decapping, 5’-3’ exonuclease, and 3’-deadenylation [84–86] (reviewed in [34, 87]). P-bodies are dynamic granules with constant exchange of many RNA-binding proteins (RBPs) and RBP components, like Pat1b and 4E-T [88–90]. While most p-body proteins have not been reported to shuttle between nuclear and cytoplasmic compartments, treatment of cells with leptomycin B or mutations that block nuclear export caused p-body-resident Pat1b and 4E-T to relocalize to the nucleus, with Pat1b localizing to NSs [89, 91, 92]. Treatment with leptomycin B also resulted in p-body loss [89], supporting the fact that RBP nucleocytoplasmic shuttling maintains p-body homeostasis. As such, events that may disrupt this homeostasis, like an abundant RNA stuck in NSs, could influence p-body dynamics despite existing in separate compartments. We showed that both WT and ΔStarts overexpression causes p-body loss, enhanced disassembly when *kaposin* NSs are highly visible (**Fig. 4B,D**) suggesting that *kaposin induces* p-body loss in a cell autonomous manner. Future work will determine if *kaposin*-induce p-body loss is a direct result of Pat1b, 4-ET, or other p-body resident RBPs getting sequestered in *kaposin* NSs.

P-bodies are enriched in ARE-mRNAs that are normally labile [93, 94] and encode for potent growth factors, pro-inflammatory cytokines, pro-angiogenic molecules, and cell cycle regulators [95, 96]. It is currently believed that p-bodies control mRNA processing by acting as mRNA storage sites, repressing transcripts until they are released for translation [84, 97, 98]. We observe this with AltB overexpression, where KapB-induced p-body loss (**Fig. 4B,D**) corresponds with an overall increase in ARE-mRNA levels (**Fig. 4E**). However, while p-body loss is classically associated with enhanced ARE-mRNA stability and expression [57, 59, 98–100], the opposite is true with ΔStarts overexpression where *kaposin*-induced p-body loss (**Fig. 4B,D**) produced an overall decrease in ARE-mRNA levels (**Fig. 4E**). Since NSs control RNA processing and splicing [42], and the formation of *kaposin* NSs alters both viral and host gene expression [38], it is possible that *kaposin* NSs also elicit changes to transcription, stability or mRNA export of cytokine transcripts that contain AREs. If true, this effect would dominate over their post-transcriptional regulation in p-bodies. Future work will assess changes in the enrichment of ARE-mRNAs within p-bodies, and their translational output, to determine if *kaposin*-mediated p-body loss is a result of transcriptional ARE-mRNA regulation.

P-bodies dynamics are also regulated by cytokine signalling. Ectopic treatment of IL-4 and TNF in bronchial epithelial cells, and treatment of IL-6 in sensory neurons, results in p-body loss [101, 102]. Conversely, treatment of mammary epithelial cells with TGF-B promoted the formation of p-bodies [103]. All of these cytokines are translated from ARE-mRNAs, suggesting a feed forward loop connecting p-body loss to the production of cytokines that then amplify p-body loss via autocrine or paracrine signalling. It was previously discovered that KapB uses its DR2 amino acid repeats to bind and activate MK2, resulting in the detectable secretion of cytokines including IL-6 [20, 21]. We found that KapB elicits p-body loss in a paracrine manner (**Fig. 5A,B**), with a corresponding overall increase in ARE-mRNAs including *IL-6* (**Fig. 4E**). It is possible that AltB expression induces cytokines secretion to antagonize p-bodies in bystander cells. Future work will test whether *kaposin* or KapB activates MK2 by monitoring its activation via the phosphorylation of downstream target hsp27 on Serine-82. As phospho-hsp27 can influence actin stress fibre formation and inhibit Hippo kinases to promote nuclear YAP translocation [104–107], future studies should test whether *kaposin* or KapB mediates hsp27 phosphorylation to uncouple these phenotypes.

### The multifunctional kaposin locus utilizes RNA and protein components to promote tumourigenesis at different phases of KSHV replication

In summary, we show that the products of the KSHV kaposin gene locus, *kaposin* and KapB, distinctly influence pro-tumourigenic pathways. Using ΔStarts overexpression as a reductionist system for latency, we show that **kaposin** RNA induces p-body loss (**Fig. 4B,D**) but fails to increase the transcription of cytokine and inflammatory genes (*Fig. 4E*). A potential consequence of this is dampening of the immune response which would otherwise warn immune cells of the latent infection. We also show that *kaposin* increases the nuclear translocation of YAP (**Fig. 2G**), perhaps just enough to promote latent cell survival (**Fig. 1E**, **Fig. 3F**). Using AltB as our reductionist system for lytic infection, we show that KapB induces actin SF formation (**Fig. 2A,B**), enlarges cells (**Fig. 2C,D**), magnifies the levels of total YAP (**Fig. 2C,E**; **Fig. 3A,B**), and promotes a pro-inflammatory and pro-angiogenic environment by inducing the transcription of related genes (**Fig. 3E**; **Fig. 4E**). High expression of these molecules can be toxic to cells [108, 109], which corroborates our data showing that KapB reduces total cell yield (**Fig. 1E**) and total cellular metabolic activity (**Fig. 3F**). The bifunctional ability of *kaposin* RNA shows that both *kaposin* and KapB manipulate pro-survival and pro-tumourigenic pathways, while the spatiotemporal regulation of *kaposin* translation into KapB mediated by NS sequestration highlights a novel strategy of viral subversion of host cell biology.

### Limitations

We acknowledge the following limitations in our study. First, we have not made a direct comparison between our lentivirus overexpression model and an infection model, yet our use of a reductionist approach to model kaposin locus gene expression in KSHV latency versus lytic reactivation is a valuable way to simplify a complicated heterogenous infection, and to attribute phenotypes to *kaposin* versus KapB. Second, our AltB construct is optimized for KapB translation but still translates KapA and does not encode for KapC. Future efforts should explore the design of a recoded panel of all Kap proteins to directly assess the individual contributions of these proteins. Third, A549 and EA.hy926 are both immortalized cell lines which may not fully capture the physiological consequences of the primary ECs within KS lesions. Fourth, our paracrine signalling assay supports a role for secretory factors in AltB overexpressing cells to modulate p-body dynamics without identification of the specific factor(s) responsible. Analysis of conditioned media components should be investigated.

## ACKNOWLEDGEMENTS

We thank Drs. Anne Vaahtokari and Luc Provencher of the Charbonneau Microscopy Facility for microscopy support. We acknowledge and thank all members of the Corcoran Lab for their insightful guidance and support throughout this study, and Madeline Day for her knowledge and expertise in cloning and site-directed mutagenesis. MPBM was supported by a CSM graduate training award, a Snyder Institute Beverley Phillips Doctoral training award and an Alberta graduate scholarship. This study was supported by operating funds awarded to JAC from the Natural Sciences and Engineering Research Council of Canada (NSERC) Discovery Grant RGPIN-2022-03854.

## AUTHOR CONTRIBUTIONS

**Maxwell P. Bui-Marinos**: Conceptualization; Data curation; Formal analysis; Investigation; Methodology; Validation; Visualization; Writing – original draft; Writing – review and editing.

**Jennifer A. Corcoran**: Conceptualization; Funding acquisition; Project administration; Resources; Supervision; Writing – original draft; Writing – review and editing.

## METHODS

### Cell culture

HEK-293T (human embryonic kidney epithelial cells [110]; ATCC), A549 (lung carcinoma epithelial cells [111]; ATCC), and EA.hy926 (somatic hybrid cell line of primary human umbilical vein endothelial cells and A549 cells, characterized as endothelial cells [47, 48, 112, 113]; ATCC) were cultured in Dulbecco’s modified Eagle medium (DMEM; ThermoFisher) supplemented with 10% heat-inactivated fetal bovine serum (FBS, ThermoFisher), and 100 U/mL penicillin, 100 μg/mL streptomycin, and 2 mM L-glutamine (PSQ, ThermoFisher), otherwise referred to as complete media. Media prepared without penicillin or streptomycin was used as antibiotic-free media, and media prepared without serum or PSQ was used as starvation media. Cells were maintained in a humidified incubator at 37°C with 5% CO_2_ and sub-cultured every 3 – 4 days using phosphate-buffered saline and Trypsin-EDTA at a 1:10 (HEK-293T and A549) or 1:8 (EA.hy926) ratio. To seed cells for experiments, cells were sub-cultured, collected in 15 mL conical tubes, then centrifuged at 500×*g* for 10 min. Cells were resuspended in complete media and cell concentration was determined with a hemocytometer and Trypan Blue exclusion dye (ThermoFisher) to identify dead cells based on membrane integrity.

### Plasmids and cloning

The expression constructs for pcDNA-kaposin (wild type), pcDNA-kaposin-ΔStarts and pcDNA-kaposin-altKapB (AltB) have been previously described [38]. AltB was FLAG-tagged (AltB-FLAG) by site-directed mutagenesis, and all constructs (wild type, ΔStarts, AltB, and AltB-FLAG) were subcloned into pLJM1-puro using BamHI and NheI restriction sites. Plasmids used in this study can be found in **Table 1**, and site-directed mutagenesis primers in **Table 2**.

**Table 1.** Plasmids used in this study.

| Plasmid | Application | Source |
| --- | --- | --- |
| pLJM1-Puro | Empty vector control | [28] |
| pcDNA-kaposin | Cloning | [38] |
| pcDNA-kaposin-Δstarts | Cloning | [38] |
| pcDNA-kaposin-altKapB | Cloning | [38] |
| pLJM1-Puro-kaposin | Overexpression | This study |
| pLJM1-Puro-kaposin-Δstarts | Overexpression | This study |
| pLJM1-Puro-kaposin-altKapB | Overexpression | This study |
| pLJM1-Puro-kaposin-altKapB-FLAG | Overexpression | This study |
| pMD2.G | Lentivirus production | Addgene #12259 |
| psPAX2 | Lentivirus Production | Addgene #12260 |

**Table 2.**
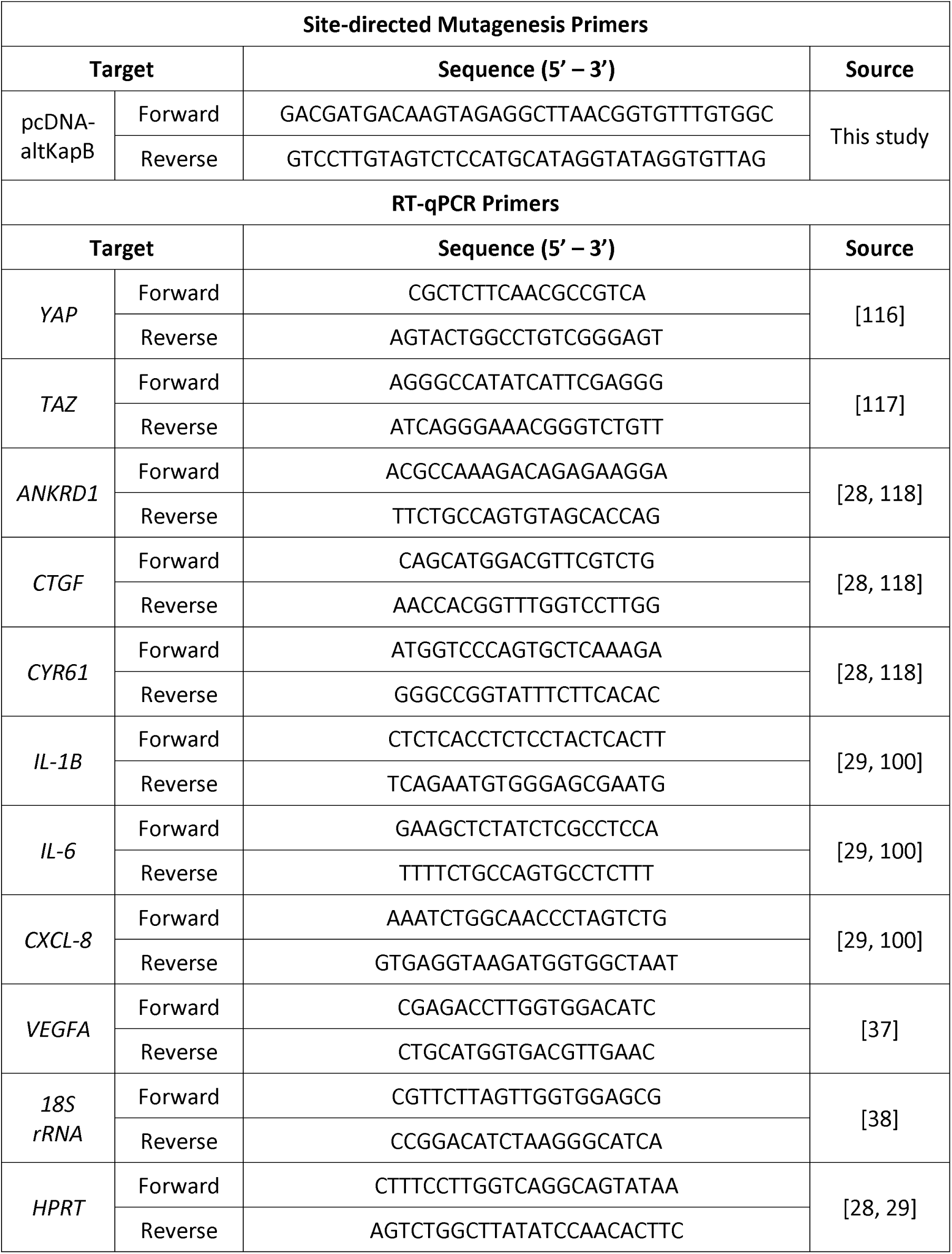
Primers used in this study.

### Recombinant lentivirus production and transduction

Second-generation recombinant lentiviruses were produced as previously described [100]. Briefly, HEK-293T cells were seeded on 10 cm dishes to achieve 70-80% confluency. The next day, transfection media was prepared by combining psPAX2, pMD2.G, and a lentiviral transfer plasmid containing the gene of interest at a 2:1:3.3 μg ratio in OptiMEM (Gibco), then mixed with polyethyleneimine (PEI, 1 mg/mL stock, Polysciences) diluted in OptiMEM to achieve 3 μL PEI/μg plasmid DNA. HEK-293T cells were washed once with PBS then treated with 4 mL of starvation media, immediately followed by the dropwise addition of 1 mL transfection media. At 6 h post-transfection, transfection supernatant was replaced with 7 mL of antibiotic-free media; after 48 h, lentivirus-containing supernatants were harvested, centrifuged at 1000×*g* for 10 mins, then filtered using a 0.45 μm polyethersulfone (PES) syringe filter (VWR). Lentivirus was aliquoted into 0.5 mL stocks then frozen at -80°C until use.

For transduction, A549 or EA.hy926 cells were seeded at 100,000 cells/well in a 12-well plate and left to adhere in a humidified, 37°C, 5% CO_2_ incubator. The next day, lentiviruses were thawed at 37°C, diluted in antibiotic-free DMEM to achieve equivalent *kaposin* transcript expression, then transferred onto target cells in the presence of 5 μg/mL polybrene (SigmaAldrich). After 24 h, media was replaced with selection media containing 1 μg/mL puromycin in complete DMEM. Selection proceeded for 48 h then cells were sub-cultured 1:3 or 1:4 into a 12-well plate, unless otherwise specified. Selection continued for another 24 hours then cells were allowed to recover for 24 – 48 h prior use for experiments. This way, the experimental endpoints in this study occurred at 5 – 6 days post-transduction unless indicated otherwise.

### Immunoflurescence RNA Fluorescence In Situ Hybridization (IF-FISH)

IF-FISH was conducted according to manufacturer’s protocol on sequential Stellaris FISH and immunofluorescence using adherent cells, with modifications. Briefly, cells seeded on coverslip were fixed at the experimental endpoint with 0.5 mL 4% (v/v) paraformaldehyde for 30 min at room temperature, then cells were permeabilized in 0.5 mL 0.1% (v/v) Triton X-100 in PBS for 10 min at room temperature. Coverslips were incubated with primary antibodies diluted in PBS overnight at 4°C. The next day, coverslips were washed with PBS to remove primary antibodies, incubated with secondary antibodies diluted in PBS for 1 h at room temperature, then fixed in 0.5 mL 4% (v/v) paraformaldehyde for 10 mins at room temperature. Coverslips were washed once with PBS then incubated in 0.5 mL Wash Buffer A (LGC Biosearch) for 5 mins. Stellaris probes for kaposin Quasar 670 have been previously reported; the same kaposin probe sequence set was used to generate another set of custom probes conjugated with fluorescein (FAM) which has an excitation wavelength at 495 nm. Stellaris probes were diluted in Hybridization Buffer (LGC Biosearch) to a concentration of 62.5 nM and coverslips were incubated in a humidified 37°C chamber overnight. The next day, coverslips were washed with 0.5 mL Wash Buffer A for 30 min, incubated with Wash Buffer A containing 5 ng/mL DAPI (Invitrogen) for 1 h at room temperature, then incubated with Wash Buffer B (LGC Biosearch) for 10 mins. Samples were mounted onto microscope slides with ProLong Gold Antifade mounting media (Thermo Fisher). To stain for F-actin, after coverslips were incubated with Wash Buffer B for 5 min, they were washed twice with 0.5 mL 2× saline-sodium citrate buffer (SSC; 300 mM sodium chloride, 300 μM sodium citrate pH 7.0) then incubated with phalloidin-conjugated AlexaFluor-647 (Invitrogen, diluted in DMSO as a 400× stock) in 2×SSC for 1.5 h at room temperature then washed once with 2×SSC prior to mounting. All confocal images were taken using a Zeiss LSM 800 Airyscan confocal microscope using the 63× 1.4 NA Oil Plan-Apo objective and the following lasers (excitation wavelength): Diode (405 nm), Argon (488 nm), and Helium/Neon (561nm and 633 nm) with emission filter ranges of 410 – 513 nm, 489 – 605nm, 570 – 659nm, and 638 – 755nm, respectively. At least two z-stacks were acquired per condition, per replicate, and collated using maximum intensity projections (MIPs) of 18 – 21 slices with 1912 × 1912 pixel dimensions. All widefield images were taken using a Zeiss AxioObserver Z1 microscope with a 40× oil-immersion objective. Antibodies used for IF-FISH are listed in **Table 3**.

**Table 3.** Antibodies used in this study.

| Antibody | Species | Application | Dilution | Source |
| --- | --- | --- | --- | --- |
| anti-Kap DRs | Rabbit | Immunoblot | 1:5000 | [38] |
|  |  | IF-FISH | 1:1000 |  |
| anti-SRMM2 | Mouse | IF-FISH | 1:200 | Abcam (ab11826) |
| anti-YAP/TAZ | Mouse | IF-FISH | 1:200 | Santa Cruz (sc-101199) |
| anti-YAP | Rabbit | Immunoblot | 1:1000 | CST (4912) |
| anti-TAZ | Rabbit | Immunoblot | 1:1000 | CST (8418) |
| anti-P-YAP/P-TAZ | Rabbit | Immunoblot | 1:1000 | CST (4911) |
| anti-DDX6 | Rabbit | IF-FISH | 1:1000 | Bethyl Labs (A300-461A) |
| Anti-FLAG | Rabbit | IF-FISH | 1:1000 | CST (14793S) |
| AlexaFluor-488<br>anti-mouse IgG | Goat | IF-FISH | 1:1000 | ThermoFisher (A-21121) |
| AlexaFluor-488<br>anti-rabbit IgG | Chicken | IF-FISH | 1:1000 | ThermoFisher (A-21441) |
| AlexaFluor-555<br>anti-rabbit IgG | Goat | IF-FISH | 1:1000 | ThermoFisher (A-11064) |
| AlexaFluor-647<br>anti-mouse IgG | Chicken | IF-FISH | 1:1000 | ThermoFisher (A-21463) |
| AlexaFluor-647<br>anti-rabbit IgG | Chicken | IF-FISH | 1:1000 | ThermoFisher (A-21443) |
| HRP anti-Rabbit IgG | Goat | Immunoblot | 1:3000 | CST (7074S) |

### Immunoblotting and densiometric analysis

Cell protein lysates were harvested in 2× Laemmli buffer (62.5 mM Tris-HCl pH 6.8, 20% glycerol, 2% SDS, 0.01% w/v bromophenol blue), chromatin sheared using a 22-gauge needle and syringe, then stored at- 20°C until use. Relative protein concentration was quantified against a bovine serum albumin (BSA; ThermoFisher) standard curve using the DC Protein Assay (BioRad) as per manufacturer’s specifications. Dithiothreitol (DTT; Sigma Aldrich) was added to protein lysates at a final concentration of 100 mM, then lysates were boiled at 95°C for 10 mins. Between 8 – 10 μg of protein was resolved by SDS-PAGE using 12% TGX Stain-Free acrylamide gels (BioRad). Proteins were transferred onto a polyvinylidene difluoride (PVDF) membrane using a Trans-Blot Turbe Transfer System (BioRad) then total protein images were acquired on a ChemiDoc Touch Imaging System (BioRad). Membranes were blocked in either 5% skim milk (anti-Kap DRs polyclonal antibody) or 5% BSA (all commercial antibodies) in TBS-T (Tris-buffered saline, 0.1% Tween-20) for 1 h at room temperature then incubated with primary antibodies overnight at 4°C. The next day, membranes were washed with TBS-T, secondary antibodies were diluted in either 5% skim milk or 5% BSA (anti-phospho-YAP/TAZ only) in TBS-T, and membranes were incubated with secondary antibody for 1 h at room temperature. Membranes were developed using Clarity Western ECL (BioRad, for overexpressed proteins) or ClarityMax Western ECL (BioRad, for endogenous protein expression) then visualized on a ChemiDoc Touch Imaging System. Densitometry analysis was conducted using BioRad ImageLab software, normalized to total protein. Antibodies used for immunoblotting are listed in **Table 3**.

### TaqMan transcript copy number analysis and reference gene RT-qPCR

RNA was harvested from cells using the RNeasy Plus Mini Kit (Qiagen) according to the manufacturer’s instructions, with the inclusion of an on-column DNase I (ThermoFisher) digestion step to remove contaminating DNA. In parallel, cells were counted from matched wells using a hemocytometer without Trypan Blue exclusion dye for transcript copy number (TCN) analysis. RNA was eluted in nuclease-free water (ThermoFisher) then stored at -80°C until further use. RNA quality was determined using a NanoDrop OneC (ThermoFisher) and concentration was determined using Qubit RNA Broad Range assay (ThermoFisher).

To assess *kaposin* transcript copy number, 500 ng of RNA was reverse transcribed into cDNA using MaximaH Minus Master Mix (ThermoFisher) in a 20 μL reaction (25 ng RNA/μL) then diluted 1:10 in nuclease-free water (represents 2.5 ng RNA/uL). Probe-based RT-qPCR was conducted using a custom TaqMan assay specific for a 164-nucleotide sequence from the KapA region of kaposin (ThermoFisher, Assay ID AP7D4GA supplied as 20× stock) as previously described [38]. Briefly, RT-qPCR actions were prepared in TaqMan Universal PCR Master Mix (ThermoFisher) as per manufacturer’s instructions, with reactions containing 1×TaqMan assay and 2.5 μL diluted cDNA meaning each reaction is estimated to represent 6.25 ng of RNA. Hemocytometer counts (cells) were divided by matched sample RNA yield (RNA yield in ng = RNA concentration × elution volume) to estimate the number of cells represented per unit RNA (cells/ng RNA). This was multiplied by the amount of RNA represented in the qPCR reaction (i.e., 6.25 ng) to estimate the number of cells represented in each reaction. The resulting reaction Cq values were related to a standard curve developed using the pcDNA-kaposin-ΔStarts plasmid, where 1 plasmid copy equals 1 *kaposin* transcript to estimate total *kaposin* transcript copy number (TCN) detected in the reaction. Reaction TCNs were divided by the number of cells represented in each reaction to approximate the number of *kaposin* transcripts detected on a per cell basis (TCN/cell) within a sample.

For reference gene analysis, 1000 ng of RNA was reverse transcribed into cDNA using MaximaH Minus Master Mix in a 20 μL reaction. cDNA was diluted 1:10 in nuclease-free water and RT-qPCR reactions contained 2.5 μL of diluted cDNA, 5 μL of 2×SsoFast EvaGreen Supermix (BioRad), and 500 nM of forward and reverse primers in a final reaction volume of 10 μL. Reactions were prepared in 96-well PCR plates (BioRad) and run on a BioRad CFX Connect. The cycling conditions were as follows: initial denaturation at 95°C for 3 mins, 40 amplification cycles of (denaturation at 95°C for 3 s, annealing at extension at 60°C for 10 s), and final extension at 60°C for 3 mins. Reactions were followed by a melt curve analysis between 60°C – 95°C increasing at 0.2°C/5 s increments. The ΔΔquantitation cycle (Cq) method was used to determine relative transcript abundance using *18S* or *HPRT* as a housekeeping gene. Primers used for RT-qPCR are listed in **Table 2**.

### Image analysis

#### Monitoring changes in actin intensity profiles

Actin intensity profiles were determined using Fiji, a plugins-included distribution of ImageJ2 for the unbiased processing and analysis of images [114]. Individual actin channel MIPs were set to grayscale, exported as a high-resolution .tif file, then opened in Fiji. The “Straight Line” tool was used to draw a line of at least 2 inches in length across cells, within the bounds of the cell edge defined by actin staining. Analyze > Plot Profile was applied to the drawn line to measure the gray value (a unitless number that corresponds to pixel intensity) along the line’s distance. Data was saved as a .csv file and graphs were generated using GraphPad Prism v10.0.

#### Estimation of cell size using total YAP/TAZ intensity

Cell size estimation and total YAP/TAZ intensity were determined using an unbiased image analysis pipeline generate in CellProfiler v4.2.8 [115] as in [28]. “IdentifyPrimaryObjects” was used to identify individual cells based on the nucleus staining channel, applying a size detection filter between 30 – 200 pixels with Global Manual thresholding. Next, “IdentifySecondaryObjects” was used on the YAP/TAZ staining channel with the Propagation method, Adaptive threshold strategy, and Robust Background thresholding method to define cell borders based on local changes in YAP/TAZ pixel intensity to decide whether a pixel is foreground or background. This identifies the cells (secondary object) for each identified nuclei (primary object). “IdentifyTertiaryObjects” was used to define the cellular cytoplasm (tertiary object) by subtracting the nucleus area from the defined cell area. The “MeasureObjectSizeShape” and “MeasureObjectIntensity” modules were applied to all objects to quantify pixel area (to approximate cell size) and total pixel intensity (to assess total YAP/TAZ intensity), respectively. Prior to data analysis, a filter was applied to remove datapoints with a nucleus pixel area < 1000, otherwise deemed to be artifacts.

#### P-body quantification

CellProfiler was used to quantify the number of DDX6-positive foci per cell as previously described [100]. Briefly, “IdentifyPrimaryObjects” and “IdentifySecondaryObjects” modules were applied as described above to identify individual cells and cell borders using the DDX6 staining channel. DDX6 foci were enhanced using the “EnhanceOrSuppressFeatures” module with a feature size of 8. The nuclei primary object was used as an inverse mask over the DDX6 channel to remove the nuclear area, thus defining DDX6 foci in the cytoplasm of cells. DDX6 foci were thresholded using an Adaptive Sauvola method with an adaptive window size of 10 to remove background staining. Individual DDX6 foci were identified using “IdentifyPrimaryObjects” with a size detection filter of 3 – 13 pixels with Adaptive Sauvola thresholding with an output function to quantify the number of DDX6 foci in individual cell cytoplasms. “MeasureObjectSizeShape” module was applied nuclei primary objects to quantify pixel area; prior to data analysis, a filter was applied to remove datapoints with a nucleus pixel area < 1000.

#### Stratifying cell populations based on high or low kaposin expression

To identify high *kaposin* expression, *kaposin* foci were identified and enhanced using the “EnhanceOrSuppressFeatures” module with a feature size of 20. The nuclei primary object was used as a mask over the kaposin channel to remove any non-nuclear kaposin signal. *Kaposin* foci were thresholded using an Adaptive Sauvola method with an adaptive window size of 5, and individual *kaposin* foci were identified using “IdentifyPrimaryObjects” applying a size detection filter of 3 – 30 pixels with Adaptive Sauvola thresholding. “MeasureObjectNeighbours” was used to measure the number of neighbouring *kaposin* foci with a distance proximity of 80 pixels. Individual *kaposin* foci (children) was merged into a single parent object based on distance proximity of 80 pixels, then parent objects were filtered based on the number of children present with a cutoff of at least 3 – 5 children and up to 100. Remaining parent kaposin foci was related to overlapping nucleus objects to identify cells with high kaposin expression. A negative mask was then applied to identify cells with low kaposin expression. This stratification was applied to measurement modules for object size and intensity, or for p-body foci quantification, to separate counts based on high or low *kaposin* expression.

### Resazurin assay

A549 cells were seeded at 10,000 cells/well in a 96-well plate and left to adhere overnight in a humidified 37°C, 5% CO_2_ incubator overnight. The next day, cells were transduced with kaposin construct recombinant lentiviruses, alongside EV and eGFP controls, in 100 μL of transduction media for 24 h. The next day, lentivirus was removed and cells were immediately recovered in 200 μL of complete DMEM. Cells were incubated in a humidified 37°C, 5% CO_2_ incubator until experimental time end points. Days 1 (post-recovery), 3, 5, and 7 post-transduction, media was removed and cells were incubated with 90 μL of complete DMEM mixed with 10 μL of Presto Blue Cell Viability Reagent (Invitrogen) for 1 h at 37°C. The fluorescence of resorufin, the metabolic product of resazurin reduction by adherent cells, is measured using a plate reader with Excitation Wavelength of 560 and Emission Wavelength of 590 nm.

### Cell-conditioned media production and paracrine signalling assay

EA.hy926 cells were seeded at 100,000 cells/well in a 12-well plate and left to adhere overnight in a humidified 37°C, 5% CO_2_ incubator overnight. The next day, cells were transduced with kaposin construct recombinant lentiviruses as described, alongside EV, eGFP and no transduction controls. Post-selection, cells were sub-cultured 1:1 into a 6-well plate and incubated in a humidified 37°C, 5% CO_2_ incubator until cells reached confluency and contact inhibition (about 48 – 72 h). To generate cell-conditioned media (CM), cells were washed 3 × 1 mL PBS then incubated with 1 mL starvation media. After 48 h, CM was collected, centrifuged at 1000 x *g* for 10 min to pellet cellular debris, and the resulting supernatant containing clarified CM was transferred to a new 1.5 mL Eppendorf and stored at -80°C until use.

To assess for paracrine signalling of CM secretory factors, naïve EA.hy926 cells were seeded onto 18mm round-bottom coverslips at 150,000 cells/coverslip and left to adhere in a humidified, 37°C, 5% CO_2_ incubator. After 48 h, cells were washed 3×1 mL coverslips at 150,000 cells/well then incubated for 48 h to adhere. Cells were washed 3 × 1 mL PBS then incubated with 0.5 mL of starvation media at 37°C to sensitize cells. After 2 h, cell supernatants was removed and cells were treated with 0.5 mL of CM or starvation media for 2 h at 37°C. Cells were then prepared for immunofluorescence.

### Immunofluorescence (IF)

Cells seeded onto coverslips were fixed at the experimental endpoint with 0.5 mL 4% (v/v) paraformaldehyde in PBS for 30 mins at room temperature. Cells were permeabilized in 0.5 mL 0.1% (v/v) Triton X-100 in PBS for 10 min at room temperature and blocked in 1% human AB serum (Sigma Aldrich) in PBS for 1 h at room temperature. Coverslips were washed with PBS then incubated with primary antibodies (anti-YAP/TAZ and anti-DDX6) diluted in 1% human AB serum solution overnight at 4°C. The next day, coverslips were washed with PBS, incubated with secondary antibody and 1 g/mL Hoechst-33342 (Invitrogen) for 1 h at room temperature, then mounted onto microscope slides with ProLong Gold Antifade mounting media. Cells were co-stained with anti-mouse AlexaFluor-488 and anti-rabbit AlexaFluor-647 to minimize bleed through. For quantification of p-bodies and assessing total YAP/TAZ intensity, images were taken using a Zeiss AxioObserver Z1 microscope with a 40× oil-immersion objective. Antibodies used for immunofluorescence are listed in **Table 3**.

### Statistics

All statistical analyses were performed using GraphPad Prism v10.0. Prior to application of statistical tests, datasets were subjected to a Shaprio-Wilk normality test to determine whether data is parametric or non-parametric. All microscopy, image-based datasets followed a non-parametric trend, and so a Kruskal-Wallis H test with Dunn’s post-hoc was applied for the multiple comparison of independent groups. All other data followed a parametric trend; either a repeated-measures one-way ANOVA with Tukey’s post-hoc or uncorrected Fisher’s least significant difference post-hoc, ordinary one-way ANOVA with Tukey’s post-hoc, or two-way repeated-measures ANOVA with a Bonferonni correction was applied to parametric datasets. Statistical tests applied are identified in respective figure legends.

